# Hemispheric asymmetries in posterior alpha power reflect the selection and inhibition of spatial context information in working memory

**DOI:** 10.1101/681031

**Authors:** Marlene Rösner, Stefan Arnau, Isabel Skiba, Edmund Wascher, Daniel Schneider

**Affiliations:** Leibniz Research Centre for Working Environment and Human Factors, Dortmund, Germany; Faculty of Psychology, Ruhr-University Bochum

**Keywords:** working memory, selective attention, neural oscillations, inhibitory control, proactive interference

## Abstract

There is an ongoing debate on the contribution of target enhancement and distractor inhibition processes to selective attention. In a working memory task, we presented to-be-memorized information in a way that posterior hemispheric asymmetries in oscillatory power could be unambiguously linked to lateral target vs. distractor processing. Alpha power asymmetries (8-14 Hz) were insensitive to the number of cued or non-cued items, supporting their relation to spatial attention. Furthermore, we found an increase in alpha power contralateral to non-cued working memory content and an alpha power suppression contralateral to relevant information. These oscillatory patterns relative to the positions of cued and non-cued items were related to the participants’ ability to control for the impact of irrelevant information on working memory retrieval. Based on these results, we propose that spatially specific modulations of posterior alpha power are related to accessing vs. inhibiting the spatial context of information stored in working memory.

## 1. Introduction

There is a dynamic and rich stream of visual information from the environment that we have to deal with simultaneously. This requires to focus on and keep track of those inputs that are behaviorally relevant and to filter out those that are not. At this point, selective attention becomes important: We are able to bias information processing in favor of task relevant information, either by enhancing mental representations of target stimuli or potentially by inhibiting inputs that are irrelevant (Desimone & Duncan, 1995). These attentional control processes can act in support of perception by proactively biasing information processing in favor of anticipated targets (Rihs, Michel, & Thut, 2009; Snyder & Foxe, 2010) or by reactively deploying attention towards stimuli identified as task-relevant (Hickey, McDonald, & Theeuwes, 2006; Moher & Egeth, 2012). In addition, attention can be retroactively deployed on the level of working memory contents, meaning the mental representations of stimuli that are held activated over the short-term in order to enable detailed analysis and categorization (Baddeley, 1996; Baddeley & Hitch, 1974; Cowan, 1999). The current investigation was designed to further clarify how attentional selection within this memory system is instantiated.

In general, attentional deployment is thought to function from two sides: Mental representations of stimuli with task-relevant features are enhanced, while irrelevant inputs might be inhibited. This leads to a relative advantage for relevant stimuli and guarantees their representation in processing instances engaged in higher-level cognition and behavioral control (Desimone & Duncan, 1995; Luck, Chelazzi, Hillyard, & Desimone, 1997). For example, research concentrated on investigating the contribution of target enhancement and distractor inhibition processes to attentional orienting in visual search (Gaspelin, Leonard, & Luck, 2015; Hickey, Di Lollo, & McDonald, 2009; Sawaki, Geng, & Luck, 2012). It could be shown that inhibitory attentional control mechanisms, reflected in the distractor positivity component (Pd) of the event-related potential of the EEG, can proactively prevent the allocation of attention to salient-but-irrelevant visual stimuli, at least when inhibitory control could be based on experience with a certain stimulus feature (Awh, Belopolsky, & Theeuwes, 2012; Wang & Theeuwes, 2018).

However, with respect to mental representations already encoded in working memory, inhibition as a cognitive process independent from target enhancement might not be a requisite mechanism: Prior investigations have shown that working memory representations that are marked as irrelevant after encoding are subject to interference by new sensory inputs (Barth & Schneider, 2018; Makovski, Sussman, & Jiang, 2008; Schneider, Barth, Getzmann, & Wascher, 2017; Schneider, Barth, & Wascher, 2017) and possibly to a rapid decay (Pertzov, Bays, Joseph, & Husain, 2013; Williams, Hong, Kang, Carlisle, & Woodman, 2013). Unattended working memory contents are thus stored in a passive and more fragile representational state (Sligte, Scholte, & Lamme, 2008; Vandenbroucke, Sligte, & Lamme, 2011), supported by the observation that they are no longer reflected in ongoing neural firing rates within cortical sites typically associated with working memory storage (Rose et al., 2016; Stokes, 2015; Wolff, Jochim, Akyurek, & Stokes, 2017). If this is the case, is there really a need for inhibitory control mechanisms in the updating of working memory contents?

In the current investigation, we argue that there is a contribution of inhibitory processes to retroactive attentional orienting. We measured posterior hemispheric asymmetries in alpha oscillations (∼8-14 Hz) relative to the position of cued vs. non-cued information in a working memory task. Prior research has shown that the orienting of attention toward lateralized working memory content is linked to a decrease in alpha power contralateral to the position of cued items (e.g., Myers, Walther, Wallis, Stokes, & Nobre, 2015; Poch, Campo, & Barnes, 2014; Poch, Capilla, Hinojosa, & Campo, 2017; Schneider, Mertes, & Wascher, 2015, 2016). First evidence for the contribution of an inhibitory process to this attentional mechanism came from investigations that included an experimental manipulation of stimulus position in a way that allowed the unambiguous association of hemispheric asymmetries in the EEG signal to the attentional processing of lateral working memory content (see also: Hickey et al., 2009; Woodman & Luck, 2003). This was done by displaying to-be-memorized stimuli above vs. below and left vs. right of central fixation (i.e., non-lateralized vs. lateralized). When cuing non-lateralized items that are equally processed within both visual hemispheres, oscillatory power in the alpha frequency range (8-14 Hz) was increased over posterior visual areas contralateral to the position of the non-cued working memory content. Furthermore, alpha power decreased contralateral to the position of a cued memory item when the non-cued item was presented on the horizontal midline (de Vries, van Driel, Karacaoglu, & Olivers, 2018; Schneider, Göddertz, Haase, Hickey, & Wascher, 2019). Importantly, also cue meaning was manipulated by cuing either the to-be-reported targets (‘remember cues’) or the irrelevant distractors (‘forget cues’). Forget cues led to an earlier onset of the inhibitory process relative to conditions with cues indicating which item to remember. The other way around, remember cues led to on earlier onset of the contralateral alpha power decrease (reflecting target selection) than forget cues that only indirectly indicated target position (Schneider et al., 2019). This orthogonal manipulation of target- and distractor-related attentional mechanisms suggested an independent contribution to the focusing of attention in working memory.

In the current study, this experimental design was combined with an approach for measuring the impact of irrelevant working memory content on behavior. This was done in order to establish a link between the neural correlates of retroactive attentional selection and working memory performance. We extended the prior experimental design by Schneider and colleagues (Schneider et al., 2019) by including varying set-sizes of the memory arrays and a control condition based on a spatially neutral retro-cue. As posterior alpha power asymmetries are typically linked to the spatial orienting of attention (Bae & Luck, 2018; Hakim, Adam, Gunseli, Awh, & Vogel, 2019), they should be independent of the number of cued vs. non-cued working memory representations. We further expected to replicate prior findings of an increase in posterior alpha power contralateral to non-cued working memory representations and a respective power decrease contralateral to the position of cued items (de Vries et al., 2018; Schneider et al., 2019). As the neutral retro-cue did not involve the attentional prioritization of either the lateralized or non-lateralized working memory content, it was used for interpreting these effects as selective target enhancement vs. distractor inhibition. We hypothesized that the increase in alpha power contralateral to the non-cued item(s) should differ reliably from the hemispheric asymmetry in the neutral control condition. This would be an indication of an inhibitory process for withdrawing the focus of attention on the level of working memory. An increased suppression of contralateral alpha power relative to the neutral condition when selectively cuing the lateral item(s) would indicate the enhanced processing at the target location.

The impact of irrelevant working memory content on performance was measured by means of different memory probe conditions following the retro-cues. Participants were instructed to compare a memory probe to the color of the cued working memory item(s) (i.e., a ‘same’ vs. ‘different’ decision). A ‘different’ response was demanded following memory probes presented in a non-cued color or a new color respective the memorized items. When presenting the memory probe in a non-cued color, this information was only recently identified as task-irrelevant and should interfere with the selective retrieval of the relevant information (Oberauer, 2005). As this effect is principally comparable to the ‘proactive interference’ of older memory content with the retrieval of currently relevant information, we will make use of this label throughout the manuscript (Monsell, 1978; Whitney, Arnett, Driver, & Budd, 2001). Prior research has shown that the ability to deal with this interference effect depends on how well cued and non-cued working memory representation can be dissociated from each other (Oberauer, 2005). We thus propose that the extent of proactive interference depends on the individual ability to focus attention on the cued information and to inhibit non-cued working memory content. Both a strong suppression of posterior alpha power contralateral to the cued position and a strong increase in alpha power contralateral to the position of non-cued information should thus counteract the proactive interference effect. These results would show how retroactive attentional mechanisms guarantee that the retrieval of information from working memory proceeds in a targe-oriented way.

## 2. Materials and Methods

### 2.1. Participants

Twenty-four participants (62.5% female, M(age) = 23.5, SD = 2.78, range = 19-30) took part in the experiment. They received 10 € per hour or course credit for participation. All participants were right-handed as indicated by means of a handedness questionnaire. None of the participants reported any known neurological or psychiatric disease and all had normal or corrected-to-normal vision. Additionally, color vision was confirmed by means of the Ishihara Test for Color Blindness. Before the beginning of the experiment, all participants gave informed consent after receiving written information about the study’s purpose and procedure. The procedure was in accordance with the Declaration of Helsinki and approved by the local ethics committee at the Leibniz Research Centre for Working Environment and Human Factors.

### 2.2. Stimuli and procedure

The stimuli were eight colored discs (RGB values: red: 255-0-0; blue: 0-0-255; green: 0-255-0; yellow: 255-255-0; magenta: 255-0-255; cyan: 0-255-255; purple: 50-0-100; orange: 205-128-0; grey: 128-128-128; average luminescence = 53.6 cd/m^2^) presented along with grey sensory filler items (144-144-144, 53.6 cd/m^2^) on a dark-grey background (38-416-33, 15 cd/m^2^). A 22-inch CRT monitor (100Hz; 1024 x 768 pixels) was used for stimulus presentation. Participants were seated with a 150 cm viewing distance to the screen.

A central fixation cross was displayed on the screen during the entire duration of the trial. At the beginning of each trial, a memory array composed of two to four items was presented for 300 ms (see figure 1). The memory items were presented on the vertical midline and on the horizontal midline. The items were presented at only one of the vertical and one of the horizontal positions (i.e., above vs. below and left vs. right of fixation). When there were two adjacent items on the horizontal or vertical positions, they were presented on hypothetical circles with 1.25° and 2.5° radius. When there was only one item on the vertical or horizontal position, it was presented on a hypothetical circle with a radius of 1.875°. The positions opposite of the presented items in each memory array were filled with grey items. These non-colored items were defined as task-irrelevant and presented to minimize the sensory disbalance of the memory array.

**Figure 1.**
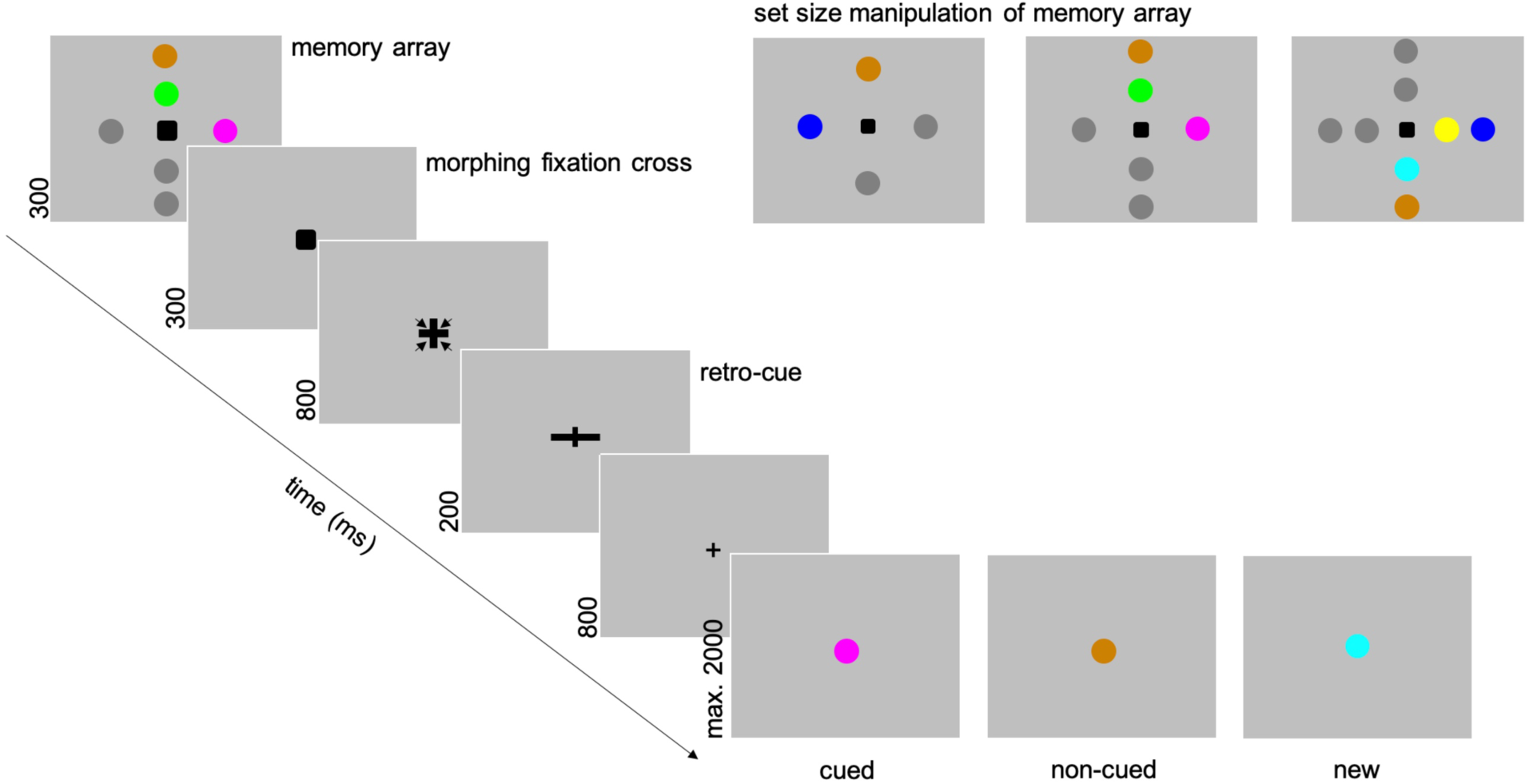
Experimental design. A memory array differing in set size from two to four relevant (colored) items was followed by a retro-cue indicating either all (neutral condition; only for set-size two) or only the lateral vs. central items as task-relevant. The later probe display either contained a stimulus in color of (one of) the cued item(s), the non-cued item(s), or was presented in a color previously not included in the memory array.

Afterwards, the fixation point started to morph into the retro-cue after a delay of another 300 ms. The morphing took 800 ms and was done in order to ensure that participants kept fixating before retro-cue onset. The cues lasted for 200 ms and indicated the position of the to-be-probed working memory items. The combinations of memory array positions and retro-cues led to conditions with either cued or non-cued lateral items (i.e., target lateral vs. distractor lateral). For memory arrays with two items, there was also a neutral retro-cue indicating both items as further on task relevant. Thus, there were overall five retro-cue conditions (set-size two with neutral cue, set-size two with one relevant and one irrelevant item, set-size three with one relevant and two irrelevant items, set-size three with two relevant and one irrelevant item and set-size four with two relevant and two irrelevant items). The number of trials within these conditions was balanced and they appeared in random order.

A probe stimulus followed the retro-cues with an inter-stimulus-interval (ISI) of 800 ms. The participants had to indicate by button press if the probe color had been presented on the memory-array position indicated by the retro-cue. There were three types of probe stimuli: cued probes (50% of all trials) when probe color had been presented on the cued position(s), non-cued or ‘recent negative’ probes (20% of all trials; condition nonexistent following the neutral cue) when probe color had been presented on the non-cued position(s) and new probes (30% of all trials) when probe color was not included in the prior memory array. The inter-trial interval varied between 500 and 1000 ms. The experiment consisted of 1440 trials that were presented in 8 blocks with 180 trials each. The blocks were separated by short breaks of around two minutes to prevent fatigue in the course of the experiment. The whole procedure took 2.5 to 3 hours, including the preparation of the EEG setup.

### 2.3. Behavioral analyses

Errors in the current experiment involved missed responses (no response within 2000 ms after probe presentation) and incorrect assignments of response categories. All responses prior to 150 ms after the onset of the memory probe were labeled as ‘premature responses’ and not included in the behavioral analyses. First, we tested for differences in error rates and response times (RTs) between the conditions including selective retro-cues. Analyses of variance (ANOVAs) with the within-subject factors *‘cued items’* (one vs. two) and *‘non-cued items’* (one vs. two) were conducted. Analyses subsequently focused on the proactive interference effect by comparing RTs and error rates between the non-cued and the new probe condition. Repeated measures ANOVAs with the factors *‘cued items’* and *‘non-cued items’* further included a within-subject factor for *probe-type* (non-cued vs. new). Additionally, error rates and RTs were compared between conditions with selective and neutral retro-cues by means of two-sided within-subject *t*-tests (only for memory arrays with two items).

### 2.4. EEG recording and preprocessing

The EEG was recorded with a 1000 Hz sampling rate from 64 Ag/AgCl passive electrodes (Easycap GmbH, Herrsching, Germany) in extended 10/20 scalp configuration. A NeurOne Tesla AC-amplifier (Bittium Biosignals Ltd, Kuopio, Finland) was used for recording while applying a 250 Hz low-pass filter. Ground electrode was set to position AFz and FCz was used as online-reference. Channel impedances was kept below 10kΩ.

Data were analyzed using MATLAB® and the EEGLAB toolbox (Delorme & Makeig, 2004). After downsampling to 500 Hz, a high-pass (0.1 Hz, 0.05 Hz cutoff, 0 to −6 dB transition window) and low-pass filter (30 Hz, 33.75 Hz cutoff, 0 to −6 dB transition window) were applied before data were subsequently re-referenced to the average of all channels. Channels with kurtosis exceeding 5 SD (M=6 channels, SD=1.6) were replaced with a spherical spline interpolation of the immediately proximal channels. In order to allow for a reliable identification of eye-movements within our data, this rejection method was not applied to the anterior lateral channels (F9, F10, AF7, AF8, AF3, AF4, Fp1, Fp2). Before segmenting data into epochs from 1000 ms before to 3600 ms after presentation of the memory array, a further high-pass filter was applied (1 Hz, 0.5 Hz cutoff, 0 to −6 dB transition window). Every second trial was selected and an automatic trial rejection procedure (threshold limit: 500 µV, probability threshold: 5 SD, Max. % of trials rejected per iteration: 5%) was applied before independent component analysis (ICA) was run. Subsequently, the IC weights were applied back to the 500 Hz data (0.1 Hz high-pass filter, 30 Hz low-pass filter and rejected channels). ADJUST (Mognon, Jovicich, Bruzzone, & Buiatti, 2010) was used for detecting and removing components labeled as eye blinks, vertical eye-movements and generic data discontinuities. Additionally, single dipoles were estimated for each IC by means of a boundary element head model (Fuchs, Kastner, Wagner, Hawes, & Ebersole, 2002). ICs with a dipole solution with more than 40% residual variance were excluded from the signal. Finally, trials with residual artifacts were rejected by means of the automatic procedure implemented in EEGLAB (threshold limit: 1000 µV, probability threshold: 5 SD, Max. % of trials rejected per iteration: 5%). These preprocessing steps led to the rejection of 327 trials on average (SD=91.36).

In an additional step, we excluded trials containing strong EEG correlates of lateral eye-movements. This was done by selecting the lateral frontal channels F9/F10 and then sliding a 100 ms time window in steps of 10 ms within an interval from memory array onset to the onset of the probe display (2400 ms later). A trial was marked for rejection, if the change in voltage from the first half to the second half of at least one of these 100 ms windows at F9 or F10 was greater than 20 µV (Adam, Robison, & Vogel, 2018; Schneider et al., 2019). This led to an additional rejection of 0 to 194 trials (M=74, SD=62.839).

### 2.5. EEG time-frequency analyses

Event-related spectral perturbation (ERSP; see Delorme & Makeig, 2004) was computed by convolving complex Morlet wavelets with each EEG data epoch, with the number of wavelet cycles in the data window increasing half as fast as the number of cycles used in the corresponding fast-fourier transformation (FFT). This led to 3-cycle wavelets at lowest frequency (i.e. 4 Hz) and 11.25-cycle wavelets at highest frequency (i.e. 30 Hz). Respective values were extracted for 200 time points and for 52 logarithmically arranged frequencies from 4 to 30 Hz.

#### 2.5.1. Hemispheric asymmetries in oscillatory power

Time-frequency analyses of hemispheric asymmetries began with the calculation of oscillatory power at electrode locations contralateral and ipsilateral to the position of cued vs. non-cued working memory contents. Lateral posterior electrodes were chosen in accordance with earlier investigations (PO7/8, PO3/4, P7/8, P5/6; see Gould, Rushworth, & Nobre, 2011; Schneider et al., 2019). The contralateral minus ipsilateral differences of the distractor lateral and target lateral conditions averaged across all set-size conditions were statistically contrasted by means of within-subject *t*-tests for each time-frequency data point. Cluster-based permutation statistics were applied for correcting for multiple comparisons. Within 1000 permutations, the conditions with lateral targets vs. distractors were randomly assigned for each dataset and within-subject *t*-tests were run for each time-frequency data point. This resulted in a 52 (frequencies) x 200 (times) x 1000 (permutations) matrix. For each permutation, the size of the largest cluster of time-frequency points with *p*-values of *p* < 0.01 was assessed. Differences between conditions in the original data were considered significant, if the size of a cluster of time-frequency points with *p*-values of *p* < 0.01 were larger than the 95^th^ percentile of the permutation-based distribution of maximum cluster sizes (see solid lines in figures 3). These analyses revealed a clear significant time-frequency cluster in alpha frequency range (8-14 Hz) following the retro-cues (450-780 ms; see figure 3A). Further ANOVAs were run based on this time-frequency range and included the within-subject factors *‘cued items’* (one vs. two), *‘non-cued items’* (one vs. two) and *‘eliciting stimulus’* (cued vs. non-cued item(s) lateral). Additionally, the same time and frequency ranges were used for testing whether posterior asymmetries in the target lateral and distractor lateral conditions differed significantly from zero (one-sided tests for contralateral enhancement vs. contralateral suppression).

**Figure 3.**
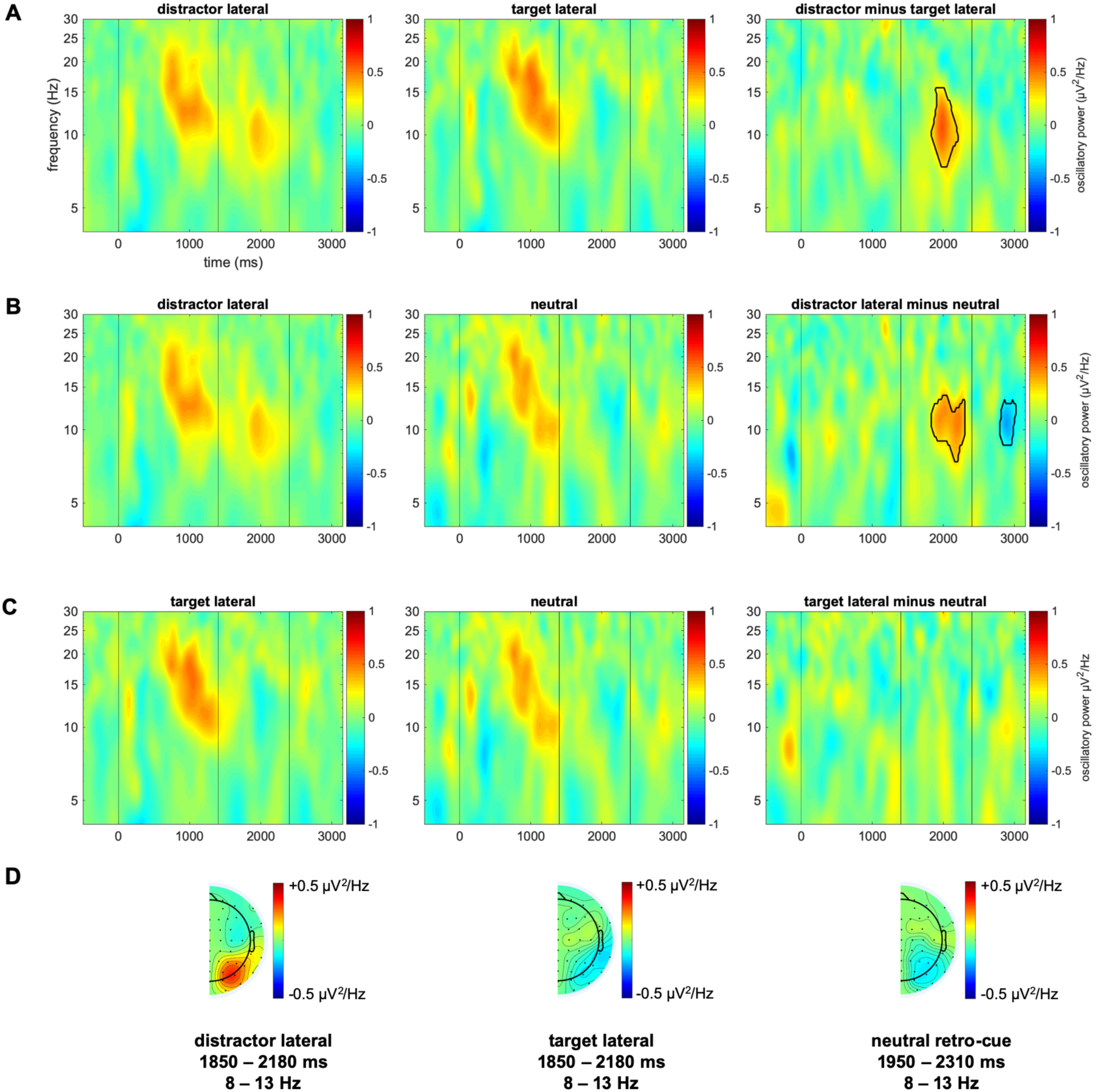
Hemispheric asymmetries in oscillatory power. 3A depicts the comparison of the distractor lateral and target lateral contralateral-ipsilateral differences. The respective comparison of the distractor lateral and neutral conditions is shown in 3B. The comparison of the target lateral and neutral condition is depicted in 3C. Vertical lines indicate the onsets of the memory array, retro-cue and memory probe. The topographies of these effects (D) are only displayed on one hemisphere, because contralateral and ipsilateral signals were averaged across left- and right hemispheres.

A comparable procedure was used for the neutral retro-cue condition. As we expected posterior asymmetries in this condition to differ from the distractor lateral condition, we compared each time-frequency point in the contralateral minus ipsilateral difference between these conditions (averaged across all memory array set-sizes for the distractor lateral condition) and then applied cluster-based permutation statistics. These analyses revealed a significant cluster from around 8 to 14 Hz and 450 to 900 ms following the cues (see figure 3B). A subsequent one-sided *t*-test was used for testing if the asymmetry in the neutral retro-cue was reliably different from zero within this time-frequency range.

Prior investigations further raised issues regarding a potential confound of hemispheric alpha power asymmetries following the retro-cues by prior lateral offsets in fixation position. This was due to the fact that already the memory array featured a lateral bias to the left or right side. We accordingly measured the contralateral vs. ipsilateral event-related potential (ERP) at frontal channels F9/F10 relative to the position of relevant lateral memory array stimuli. Already the memory array caused a contralateral negativity at F9/F10 with a peak in the grand average difference wave at 610 ms (averaged across conditions). Mean amplitudes per condition were measured within a 100 ms time window centered on this peak and within the 200 ms interval prior to retro-cue presentation. To assess if these lateral offsets in fixation prior to cue presentation might be related to the hypothesized alpha power asymmetries, we made use of repeated measures ANOVAs with *eliciting stimulus* and the number of cued and non-cued items as within-subject factors. Additionally, we included a covariate for lateral eye movements (measured as frontal ERP asymmetry in the two time windows across experimental conditions; see also: supplementary material in Schneider et al., 2019).

For all statistical analyses (behavioral and EEG data), Greenhouse-Geisser correction (indicated by Greenhouse-Geisser ε) was applied when sphericity of the data was violated. Partial eta squared (*η^2^_p_*) was used as an indicator of effect size for all ANOVAs. For post-hoc analyses, the false discovery rate procedure as indicated by Cramer and colleagues (Cramer et al., 2016) was used for correcting for cumulation of Type 1 error within the ANOVAs. In these cases, adjusted critical *p-*values (*p_crit_*) are provided. Cohen’s d_z_ was used as a measure of effect size for within-subject *t*-tests. The false discovery rate (FDR) procedure was used when post-hoc comparisons required correcting for cumulative Type 1 error (indicated as adjusted *p*-values or *p_adj_* for *t*-test parameters).

#### 2.5.2. Correlational analyses

We furthermore investigated if and to what extent the electrophysiological correlates of retroactive attentional orienting were related to the participants’ ability to control for the interference of irrelevant mental representations on working memory retrieval processes (i.e., proactive interference). Proactive interference was measured as the difference in RTs between the non-cued and new probe conditions, averaged across retro-cue conditions. As a correlate of the processing of relevant lateral working memory content, we averaged across all trials containing a lateral target. Also the neutral retro-cue condition was added to this average, as it included a relevant lateral item (and did not differ from the target lateral conditions in terms of hemispheric alpha power asymmetries, see results section). The contralateral increase in posterior alpha power relative to the non-cued positions (with targets presented on the horizontal midline) was used as a correlate of distractor inhibition. The time and frequency ranges for assessing these effects were based on the results of the cluster-permutation statistics described above (i.e., 8-14 Hz and 450-900 ms after retro-cue presentation). The individual level of posterior alpha power contralateral to relevant or irrelevant working memory content was correlated separately with the proactive interference effect on RT level (i.e., a between-subjects approach, based on Pearson correlations). Additionally, we defined an ‘attentional selectivity index’ (short: ASI) by subtracting the alpha power asymmetry for relevant lateral items from the asymmetry observed when lateral items were indicated as irrelevant. All three correlations (distractor lateral, target lateral and ASI) were statistically assessed by 10000 bootstrapped samples (i.e., random sampling with replacement) and the calculation of the 95% confidence intervals in the resulting distribution. This approach reduces sensitivity to outlier values. The *p*-values for assessing whether these correlations differ significantly from zero can be calculated by dividing the number of bootstrap instances with negative or positive *r*-values by the overall number of bootstrap instances (10000). For the distractor lateral condition and the ASI, we expected a negative correlation with the proactive interference effect (i.e., larger values indicate stronger inhibition/attentional selectivity and should lead to lower proactive interference). The other way around, a stronger contralateral decrease in alpha power as a correlate of target processing should be related to a lower interference effect on RTs (i.e., a positive correlation). FDR correction was applied to control for multiple testing.

## 3. Results

### 3.1. Behavioral data

Error rates varied with the number of cued items, with lower rates when only one item was cued, *F*(1,23)=48.169, *p*<0.001, *p_crit_*=0.033, *η^2^_p_*=0.677, and with the number of non-cued items, with lower error rates for one non-cued mental representation, *F*(1,23)=48.701, *p*<0.001, *p_crit_*=0.05, *η^2^_p_*=0.679. Additionally, the *cued x non-cued* interaction was significant, *F*(1,23)=24.367, *p*<0.001, *p_crit_*=0.017, *η^2^_p_*=0.514. Post-hoc analyses showed that error rates were lower when one of two items compared to one of three items was cued, *t*(23)=-3.250, *p_adj_*=0.004, *d_z_*=-0.663. This difference in error rates was larger when comparing the conditions with two cued items as a function of the number of non-cued items, *t*(23)=-7.804, *p_adj_*<0.001, *d_z_*=-1.593 (see figure 2). However, selective retro-cues following two-item memory arrays did not significantly reduce error rates relative to the neutral cue condition, *t*(23)=-0.893, *p*=0.381, *d_z_*=-0.182.

**Figure 2.**
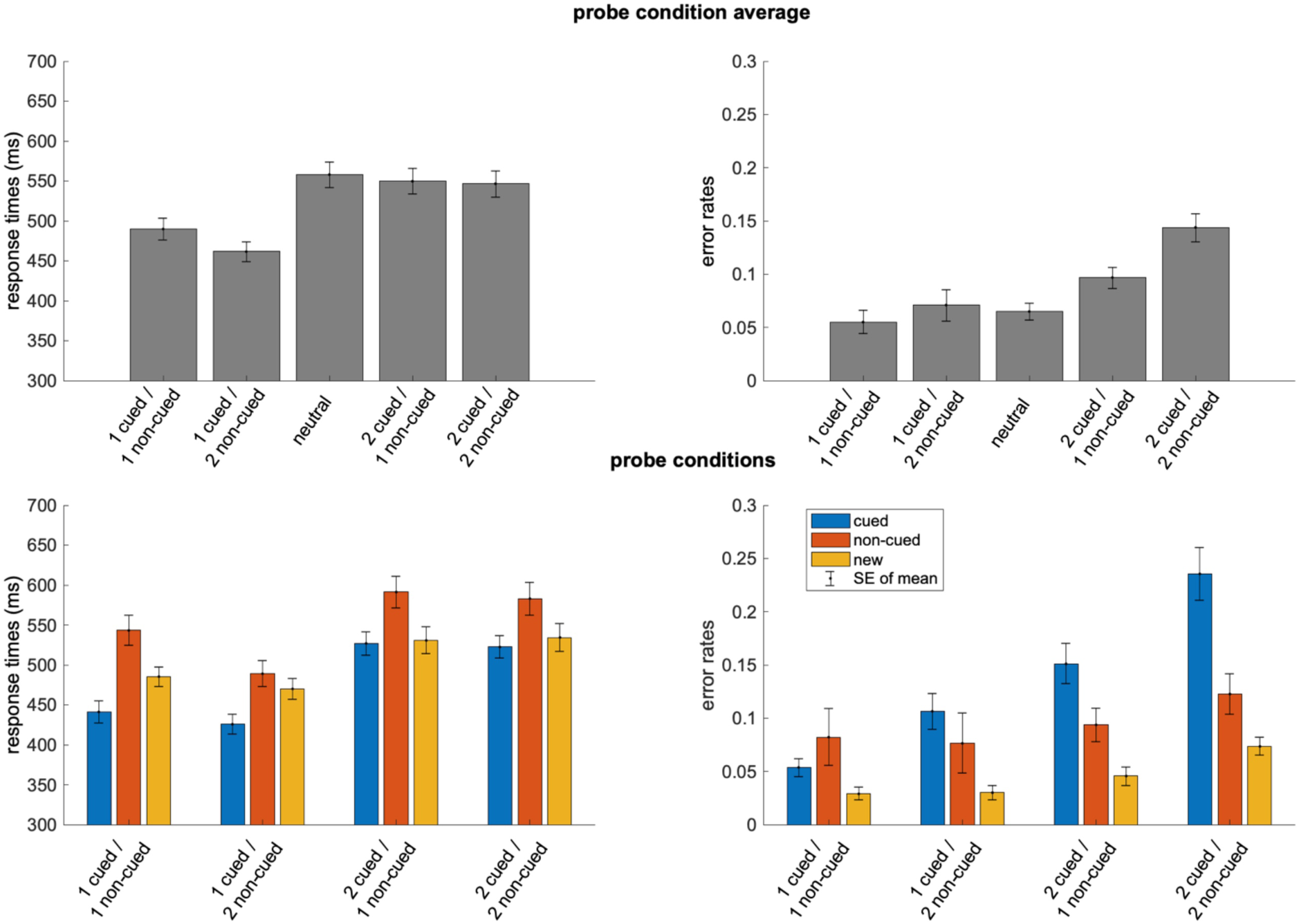
Behavioral results. The upper plots show the response times (RTs) and error rates for the different retro-cue conditions, averaged across memory probe conditions. The lower plots depict response time and error rate patterns as a function of probe condition. The neutral condition was not displayed in this regard, as it did not include a non-cued probe condition.

Further analyses concentrated on the comparison of the non-cued and new probe conditions. The non-cued probe condition featured generally higher error rates, *F*(1,23)=7.613, *p*=0.011, *η^2^_p_*=0.249. This effect was, however, not modulated by the number of cued or non-cued items (all *F*-values < 1).

RTs also varied with the number of cued items, *F*(1,23)=97.035, *p*<0.001, *p_crit_*=0.05, *η^2^_p_*=0.808. Furthermore, RTs were faster with two non-cued items compared to one non-cued item, *F*(1,23)=19.604, *p*<0.001, *p_crit_*=0.017, *η^2^_p_*=0.460, and there was a significant *cued x non-cued* interaction, *F*(1,23)=20.014, *p*<0.001, *p_crit_*=0.033, *η^2^_p_*=0.465. Post-hoc *t*-tests indicated that RTs were faster when one of three items (1 cued / 2 non-cued) compared to one of two items (1 cued / 1 non-cued) was retroactively cued, *t*(23)=5.044, *p_adj_*<0.001, *d_z_*=1.030, while no such difference as a function of the number of non-cued items was evident within the conditions based on a two-item retro-cue, *t*(23)=0.983, *p_adj_*=0.336, *d_z_*=0.201. When comparing RTs between the non-cued and new probe conditions that both demanded the same type of response, the RT difference was modulated by the number of non-cued items, *F*(1,23)=17.381, *p*<0.001, *p_crit_*=0.021, *η^2^_p_*=0.430, with a reduced RT difference for two non-cued items (cued items: *F*(1,23)=6.918, *p*=0.014, *p_crit_*=0.014, *η^2^_p_*=0.231). The *cued x non-cued x probe-type* three-way interaction was not significant, *F*(1,23)=3.951, *p*=0.059, *p_crit_*=0.007, *η^2^_p_*=0.147 (see figure 2). Additionally, RTs were faster following a selective retro-cue (only two-item memory arrays) compared to a neutral cue, *t*(23)=-7.149, *p*<0.001, *d_z_*=-1.459, indicating a general retro-cue benefit on RT level.

### 3.2. Hemispheric asymmetries in oscillatory power

As illustrated in figure 3, cluster-corrected comparisons between the target lateral and distractor lateral conditions (averaged across all conditions with selective retro-cues) revealed a latency interval with a reliable difference in alpha frequency range (8-14 Hz) from approximately 450 to 780 ms after retro-cue onset (see figure 3A). Follow-up analyses were focused on this time and frequency interval. As already indicated by the cluster-corrected *t*-tests, there was a main effect of eliciting stimulus, *F*(1,23)=33.157, *p*<0.001, *η^2^_p_*=0.590. This effect was composed of an increase in alpha power contralateral to the non-cued items, *t*(23)=6.837, *p_adj_*<0.001, *d_z_*=1.396 (*t*-test against zero), and a contralateral alpha power suppression when cued items were presented at the lateral positions, *t*(23)=-2.341, *p_adj_*=0.014, *d_z_*=-0.478 (*t*-test against zero). As hypothesized, this effect was not modulated by the number of cued items, *F*(1,23)=2.877, *p*=0.103, *η^2^_p_*=0.111, or non-cued items, *F*(1,23)=0.307, *p*=0.585, *η^2^* =0.013. Also the three-way interaction was statistically non-significant, *F*(1,23)=0.932, *p*=0.344, *η^2^* =0.039.

In a further step, we compared the neutral condition with the distractor lateral condition (averaged across memory array set-sizes) based on the cluster-correction procedure (see figure 3B). This again brought up a time-frequency area with a statistically reliable difference between conditions from 8 to 14 Hz and from 550 to 910 ms following the cues. When testing for a posterior asymmetry in alpha power based on these parameters, a contralateral suppression of alpha power in the neutral retro-cue condition was evident, *t*(23)=-1.999, *p*=0.029, *d_z_*=-0.408 (*t*-test against zero). No statistically reliable difference was observed between the target lateral condition and the neutral condition (see figure 3C).

We furthermore assessed whether lateral eye movements following memory array and prior to retro-cue presentation had any influence on the difference of alpha power asymmetries between conditions with lateral targets vs. lateral distractors. In line with earlier findings objecting such a relationship (de Vries et al., 2018; Schneider et al., 2019), the ANCOVA with the mean contralateral-ipsilateral difference from 560 to 660 ms at F9/F10 as covariate still revealed a reliable difference in posterior alpha asymmetry following lateral targets vs. distractors, *F*(1,22)=28.298, *p*<0.001, *η^2^_p_*=0.563. The interaction of *eliciting stimulus* and the eye-movement covariate was non-significant, *F*(1,22)=2.213, *p*=0.151, *η^2^_p_*=0.091. The same results were found when measuring lateral eye-movements in the 200 ms interval prior to retro-cue onset, with a statistically significant effect of *eliciting stimulus* on alpha power asymmetries, *F*(1,22)=33.604, *p*<0.001, *η^2^_p_*=0.604, and a statistically non-reliable interaction, *F*(1,22)=2.294, *p*=0.144, *η^2^_p_*=0.094. This clearly shows that the asymmetric alpha modulations following the retro-cues cannot have been caused by prior systematic effects of lateral eye movements.

### 3.3. Correlational analyses

These analyses were run in order to relate the oscillatory correlates of retroactive attentional orienting to the extent of proactive interference by non-cued working memory representations. Based on our experimental manipulations, it was possible to independently assess the contribution of target processing vs. distractor inhibition mechanisms to the handling of proactive interference. The latter was assessed as the difference in RTs between the non-cued and new probe conditions and reflects to what extent non-cued mental representations interfere with target-oriented information processing following the retro-cues. The extent of the posterior increase in alpha power contralateral to the non-cued position was negatively correlated to the non-cued minus new probe difference on RT level (r=-0.327, p=0.044). Conversely, participants with a stronger suppression of alpha power contralateral to the position of relevant working memory items revealed a reduced proactive interference effect on RT level (i.e., a positive correlation). These two correlations differed reliably in terms of the direction of the relationship between the variables of interest, as indicated by the fact that their 95% confidence intervals did not overlap. When combining these two parameters by subtracting the suppression of alpha power contralateral to relevant items from the increase in alpha power contralateral to non-cued information (labeled the ‘attentional selectivity index’ or ASI), there was also a statistically reliable correlation with the proactive interference effect on RT level. Thus, participants that could effectively bias the stored working memory representations in favor of relevant content, by focusing on the target or by de-focusing distracting information (or both), were more efficient in counteracting losses of performance related to proactive interference.

## 4. Discussion

This study investigated if and how target selection and distractor inhibition processes are engaged during the attentional prioritization of information in working memory. This was done based on a paradigm that required dealing with the interference of non-cued working memory contents during retrieval and memory probe processing. In line with earlier findings (Barth & Schneider, 2018; Makovski et al., 2008; Schneider, Barth, Getzmann, et al., 2017), behavioral results showed a performance benefit for a one-item retroactive focus of attention (compared to two-item retro-cues) and a general retro-cue benefit compared to a neutral cue condition on RT level (Griffin & Nobre, 2003; Myers et al., 2015; Schneider et al., 2015). Results further indicated that responding to a recently non-cued probe color entailed dealing with proactive interference, as both RTs and error rates were increased in this condition relative to a memory probe with a color not recently stored in working memory (see also: Oberauer, 2005; Schneider et al., 2015, 2016). Furthermore, the proactive interference effect on RT level was reduced when two items compared to one item were retroactively cued. This might indicate that the extent of proactive interference during working memory retrieval is based on the individual representational strength of the non-cued color values, that should be lower for two adjacent non-cued items compared to a single item.

The selective processing of working memory content following retroactive cues has been associated with modulations of posterior alpha power (Myers et al., 2015; Poch et al., 2014; Schneider et al., 2015, 2016). Here, we measured posterior hemispheric asymmetries relative to the cued vs. non-cued lateral positions (see figure 3). These asymmetries did not differ as a function of the number of cued or non-cued items (see also: Poch, Carretie, & Campo, 2017), in line with the notion that they reflect a spatial attentional mechanism. Importantly, we further corroborate the notion that the shifting of the focus of attention in working memory is based on inhibition: As indicated in figure 3A, hemispheric asymmetries in alpha power differed between conditions with lateral targets and lateral distractors. There was a suppression of posterior alpha power contralateral to the position of selectively cued items. This alpha power response did not differ from the neutral condition (both items cued; see figure 3C). Thus, while the posterior alpha lateralization was related to the processing of the relevant lateral item(s) in both conditions, there was no selective enhancement of processing at the cued positions. Furthermore, inhibition on the level of working memory can be conceptualized as a cognitive process that leads to a deterioration of the mental representations of already encoded but irrelevant information and proceeds independent from target selection. Our data meet these criteria, as alpha power was increased contralateral to non-cued working memory content compared to the ipsilateral oscillatory activity reflecting the processing of the opposite task-irrelevant and non-colored stimuli. While there was no sign of a selective attentional bias toward the cued items, this increase in alpha power contralateral to the non-cued positions differed reliably from the hemispheric asymmetry following neutral retro-cues (see figure 3B).

Previous findings on the contribution of inhibitory mechanisms to selective attention and their relation to posterior alpha power modulations are rather ambiguous. Some current approaches do not see sufficient evidence for a contribution of distractor inhibition (for review: Foster & Awh, 2018). Others support the functional relevance of inhibitory processes for selective information processing. On the neural level, it has been shown that increased alpha power is associated with decreased neuronal firing rates (Haegens, Nacher, Luna, Romo, & Jensen, 2011). A relation between alpha oscillations and inhibition in a psychological sense was indicated by studies revealing increased alpha power over visual areas in anticipation of distracting stimuli during a working memory task (Bonnefond & Jensen, 2012). Furthermore, Händel, Haarmeier and Jensen (2011) revealed a positive correlation between the posterior alpha power lateralization following a spatial cue and the subsequent perceptual sensitivity to information presented at the non-cued locations. However, it also has to be noted that previous findings on the focusing of attention in perceptual scenes do not necessarily have to be comparable to the attentional selection and inhibition of already encoded information in working memory.

In support of a functional relevance of the observed alpha power modulations for working memory performance, posterior alpha power asymmetries correlated with the RT differences between memory probes presented in a non-cued vs. new color. Participants with high alpha power contralateral to non-cued positions and low alpha power contralateral to the position of relevant items revealed a reduced impact of irrelevant working memory content on probe processing (i.e., a lower proactive interference effect). The most important question in this regard concerns the way in which these modulations of posterior alpha power, although typically related to spatial attentional mechanisms (e.g., Bae & Luck, 2018), can influence the proactive interference on the level of non-spatial stimulus features (here: colors). Prior research has indicated that stronger lateralization effects were found when participants oriented attention toward the position of memorized information compared to a position in empty space (Hakim, Adam, Gunseli, Awh, & Vogel, 2019). This highlights that alpha oscillations in retinotopic visual areas reflect a spatial attention mechanism that interacts with non-spatial features stored in working memory. The potentially underlying mechanism might be the linking of non-spatial stimulus features like color, shape or orientation to their spatial context. This spatial context is required for dissociating cued from non-cued items during working memory retrieval (Oberauer, 2005). The contralateral increase in alpha power relative to distractor position might thus reflect an inhibitory mechanism for unbinding the non-cued color(s) from their associated spatial position (left, right, top or bottom). The other way around, a suppression of alpha power contralateral to the position of task-relevant information might reflect the binding to its spatial context. The fact that we did not observe a difference in hemispheric alpha power asymmetries between the target lateral and neutral retro-cue conditions (see figure 3C) indicates that this spatial process can proceed simultaneously (without costs) for at least two separate locations. Both inhibiting irrelevant spatial context information and updating the binding of relevant features to their locations could guarantee an effective and target-oriented retrieval process, thereby decreasing the individual susceptibility to interference by non-cued working memory items.

In summary, we investigated the contribution of excitatory vs. inhibitory processes to the retroactive orienting of attention on the level of working memory. Working memory content retroactively cued as irrelevant still interfered with selective retrieval following a memory probe, as indicated by both error rates and RTs (i.e., proactive interference). Hemispheric asymmetries in alpha power were not modulated by the number of cued or non-cued items. However, we showed that these retinotopic alpha power modulations (see figure 3) correlated with the slowing of RTs for memory probes presented in a non-cued color (see figure 4). We argue that posterior alpha power asymmetries following retro-cues reflect the binding of relevant information to its spatial context, as well as the unbinding of irrelevant working memory content from associated locations. High efficiency in these processes guarantees a target-oriented retrieval process, thereby reducing the interference by irrelevant working memory content.

**Figure 4.**
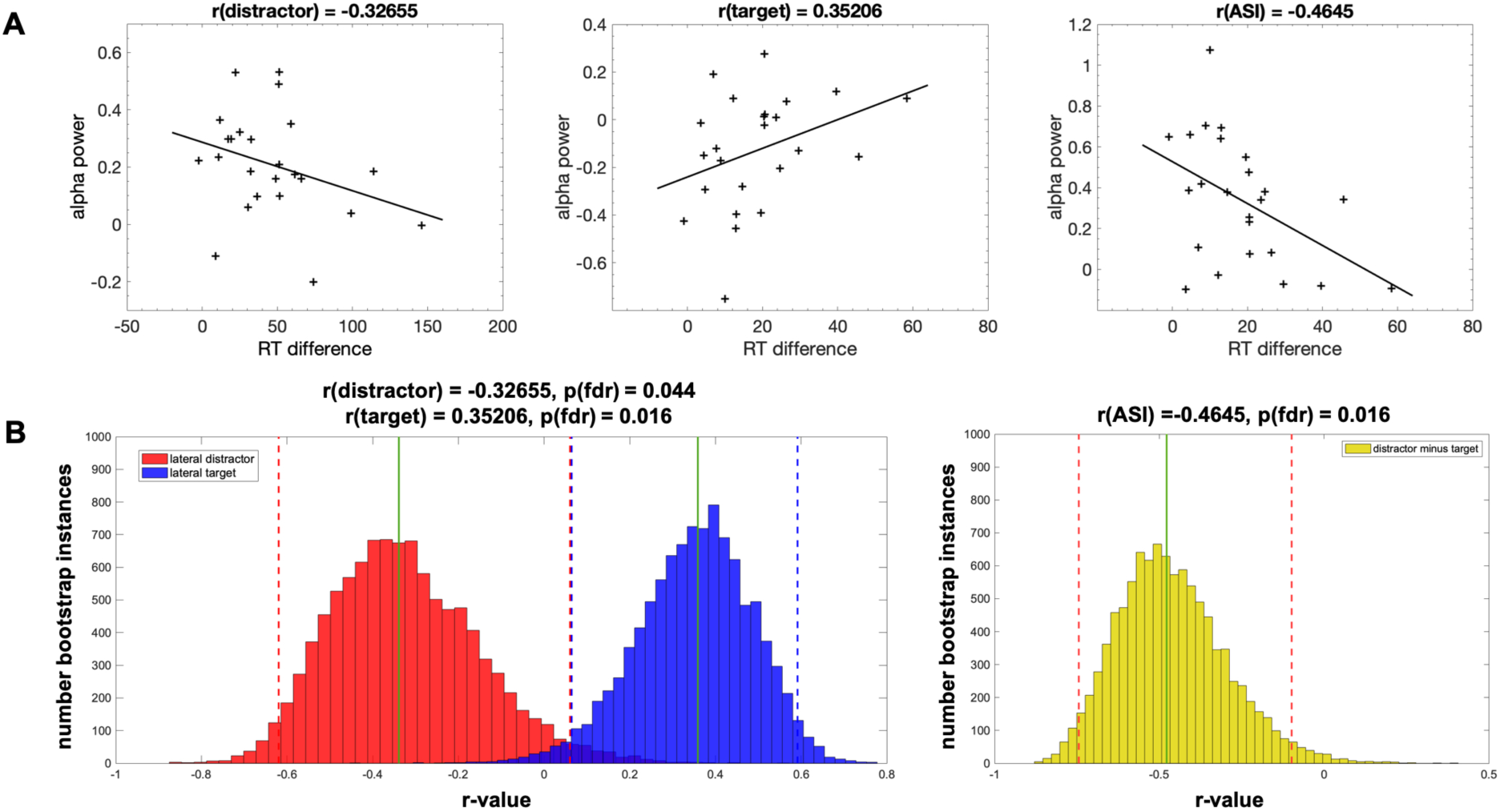
EEG-behavior correlations. The correlations were calculated for hemispheric alpha asymmetries in the distractor lateral condition, for conditions with relevant lateral items (including the target lateral and neutral conditions) and for the difference between these two values (also labeled ‘attentional selectivity index’ or ASI; see 4A). Correlations were statistically assessed based on 10000 bootstrap instances (see bar plots in 4B). A correlation was considered as statistically significant when 95% of the resulting distributions were above (for lateral targets) or below zero (for lateral distractors and ASI).

## Acknowledgements

We would like to thank Tobias Blanke for programming the experiment.

## 5. Competing interests

The corresponding author declares that financial or non-financial competing interests do not exist.

## References

Adam, K. C. S., Robison, M. K., & Vogel, E. K. (2018). Contralateral Delay Activity Tracks Fluctuations in Working Memory Performance. Journal of Cognitive Neuroscience, 30(9), 1229–1240. doi:10.1162/jocn_a_01233

Awh, E., Belopolsky, A. V., & Theeuwes, J. (2012). Top-down versus bottom-up attentional control: a failed theoretical dichotomy. Trends Cogn Sci, 16(8), 437–443. doi:10.1016/j.tics.2012.06.010

Baddeley, A. D. (1996). Exploring the central executive. The Quarterly Journal of Experimental Psychology Section A: Human Experimental Psychology, 49(1), 5–28.

Baddeley, A. D., & Hitch, G. (1974). Working Memory. In G. H. Bower (Ed.), The psychology of learning and motivation: Advances in research and theory (Vol. 8, pp. 47–89). New York: Academic Press.

Bae, G. Y., & Luck, S. J. (2018). Dissociable Decoding of Spatial Attention and Working Memory from EEG Oscillations and Sustained Potentials. J Neurosci, 38(2), 409–422. doi:10.1523/JNEUROSCI.2860-17.2017

Barth, A., & Schneider, D. (2018). Manipulating the focus of attention in working memory: Evidence for a protection of multiple items against perceptual interference. Psychophysiology. doi:10.1111/psyp.13062

Bonnefond, M., & Jensen, O. (2012). Alpha oscillations serve to protect working memory maintenance against anticipated distracters. Curr Biol, 22(20), 1969–1974. doi:10.1016/j.cub.2012.08.029

Cowan, N. (1999). An embedded-processes model of working memory. In A. Miyake & P. Shah (Eds.), Models of working memory: Mechanisms of active maintenance and executive control (pp. 62–101). New York, NY, USA: Cambridge University Press.

de Vries, I. E. J., van Driel, J., Karacaoglu, M., & Olivers, C. N. L. (2018). Priority Switches in Visual Working Memory are Supported by Frontal Delta and Posterior Alpha Interactions. Cereb Cortex, 28(11), 4090–4104. doi:10.1093/cercor/bhy223

Delorme, A., & Makeig, S. (2004). EEGLAB: an open source toolbox for analysis of single-trial EEG dynamics including independent component analysis. J Neurosci Methods, 134(1), 9–21. doi:10.1016/j.jneumeth.2003.10.009

Desimone, R., & Duncan, J. (1995). Neural mechanisms of selective visual attention. Annu Rev Neurosci, 18, 193–222. doi:10.1146/annurev.ne.18.030195.001205

Foster, J. J., & Awh, E. (2018). The role of alpha oscillations in spatial attention: limited evidence for a suppression account. Curr Opin Psychol, 29, 34–40. doi:10.1016/j.copsyc.2018.11.001

Fuchs, M., Kastner, J., Wagner, M., Hawes, S., & Ebersole, J. S. (2002). A standardized boundary element method volume conductor model. Clin Neurophysiol, 113(5), 702–712.

Gaspelin, N., Leonard, C. J., & Luck, S. J. (2015). Direct Evidence for Active Suppression of Salient-but-Irrelevant Sensory Inputs. Psychol Sci, 26(11), 1740–1750. doi:10.1177/0956797615597913

Gould, I. C., Rushworth, M. F., & Nobre, A. C. (2011). Indexing the graded allocation of visuospatial attention using anticipatory alpha oscillations. J Neurophysiol, 105(3), 1318–1326. doi:10.1152/jn.00653.2010

Griffin, I. C., & Nobre, A. C. (2003). Orienting attention to locations in internal representations. J Cogn Neurosci, 15(8), 1176–1194. doi:10.1162/089892903322598139

Haegens, S., Nacher, V., Luna, R., Romo, R., & Jensen, O. (2011). alpha-Oscillations in the monkey sensorimotor network influence discrimination performance by rhythmical inhibition of neuronal spiking. Proc Natl Acad Sci U S A, 108(48), 19377–19382. doi:10.1073/pnas.1117190108

Hakim, N., Adam, K. C. S., Gunseli, E., Awh, E., & Vogel, E. K. (2019). Dissecting the Neural Focus of Attention Reveals Distinct Processes for Spatial Attention and Object-Based Storage in Visual Working Memory. Psychol Sci, 30(4), 526–540. doi:10.1177/0956797619830384

Händel, B. F., Haarmeier, T., & Jensen, O. (2011). Alpha oscillations correlate with the successful inhibition of unattended stimuli. J Cogn Neurosci, 23(9), 2494–2502. doi:10.1162/jocn.2010.21557

Hickey, C., Di Lollo, V., & McDonald, J. J. (2009). Electrophysiological indices of target and distractor processing in visual search. J Cogn Neurosci, 21(4), 760–775. doi:10.1162/jocn.2009.21039

Hickey, C., McDonald, J. J., & Theeuwes, J. (2006). Electrophysiological evidence of the capture of visual attention. J Cogn Neurosci, 18(4), 604–613. doi:10.1162/jocn.2006.18.4.604

Luck, S. J., Chelazzi, L., Hillyard, S. A., & Desimone, R. (1997). Neural mechanisms of spatial selective attention in areas V1, V2, and V4 of macaque visual cortex. J Neurophysiol, 77(1), 24–42. doi:10.1152/jn.1997.77.1.24

Makovski, T., Sussman, R., & Jiang, Y. V. (2008). Orienting attention in visual working memory reduces interference from memory probes. J Exp Psychol Learn Mem Cogn, 34(2), 369–380. doi:10.1037/0278-7393.34.2.369

Mognon, A., Jovicich, J., Bruzzone, L., & Buiatti, M. (2010). ADJUST: An automatic EEG artifact detector based on the joint use of spatial and temporal features. Psychophysiology. doi:10.1111/j.1469-8986.2010.01061.x

Moher, J., & Egeth, H. E. (2012). The ignoring paradox: cueing distractor features leads first to selection, then to inhibition of to-be-ignored items. Atten Percept Psychophys, 74(8), 1590–1605. doi:10.3758/s13414-012-0358-0

Monsell, S. (1978). Recency, Immediate Recognition Memory and Reaction Time. Cognitive Psychology, 10, 465–501.

Myers, N. E., Walther, L., Wallis, G., Stokes, M. G., & Nobre, A. C. (2015). Temporal dynamics of attention during encoding versus maintenance of working memory: complementary views from event-related potentials and alpha-band oscillations. J Cogn Neurosci, 27(3), 492–508. doi:10.1162/jocn_a_00727

Oberauer, K. (2005). Binding and inhibition in working memory: individual and age differences in short-term recognition. J Exp Psychol Gen, 134(3), 368–387. doi:10.1037/0096-3445.134.3.368

Pertzov, Y., Bays, P. M., Joseph, S., & Husain, M. (2013). Rapid forgetting prevented by retrospective attention cues. J Exp Psychol Hum Percept Perform, 39(5), 1224–1231. doi:10.1037/a0030947

Poch, C., Campo, P., & Barnes, G. R. (2014). Modulation of alpha and gamma oscillations related to retrospectively orienting attention within working memory. Eur J Neurosci, 40(2), 2399–2405. doi:10.1111/ejn.12589

Poch, C., Capilla, A., Hinojosa, J. A., & Campo, P. (2017). Selection within working memory based on a color retro-cue modulates alpha oscillations. Neuropsychologia, 106, 133–137. doi:10.1016/j.neuropsychologia.2017.09.027

Poch, C., Carretie, L., & Campo, P. (2017). A dual mechanism underlying alpha lateralization in attentional orienting to mental representation. Biol Psychol, 128, 63–70. doi:10.1016/j.biopsycho.2017.07.015

Rihs, T. A., Michel, C. M., & Thut, G. (2009). A bias for posterior alpha-band power suppression versus enhancement during shifting versus maintenance of spatial attention. Neuroimage, 44(1), 190–199. doi:10.1016/j.neuroimage.2008.08.022

Rose, N. S., LaRocque, J. J., Riggall, A. C., Gosseries, O., Starrett, M. J., Meyering, E. E., & Postle, B. R. (2016). Reactivation of latent working memories with transcranial magnetic stimulation. Science, 354(6316), 1136–1139. doi:10.1126/science.aah7011

Sawaki, R., Geng, J. J., & Luck, S. J. (2012). A common neural mechanism for preventing and terminating the allocation of attention. J Neurosci, 32(31), 10725–10736. doi:10.1523/JNEUROSCI.1864-12.2012

Schneider, D., Barth, A., Getzmann, S., & Wascher, E. (2017). On the neural mechanisms underlying the protective function of retroactive cuing against perceptual interference: Evidence by event-related potentials of the EEG. Biol Psychol, 124, 47–56. doi:10.1016/j.biopsycho.2017.01.006

Schneider, D., Barth, A., & Wascher, E. (2017). On the contribution of motor planning to the retroactive cuing benefit in working memory: Evidence by mu and beta oscillatory activity in the EEG. Neuroimage, 162, 73–85. doi:10.1016/j.neuroimage.2017.08.057

Schneider, D., Göddertz, A., Haase, H., Hickey, C., & Wascher, E. (2019). Hemispheric asymmetries in EEG alpha oscillations indicate active inhibition during attentional orienting within working memory. Behav Brain Res, 359, 38–46. doi:10.1016/j.bbr.2018.10.020

Schneider, D., Mertes, C., & Wascher, E. (2015). On the fate of non-cued mental representations in visuo-spatial working memory: Evidence by a retro-cuing paradigm. Behav Brain Res, 293, 114–124. doi:10.1016/j.bbr.2015.07.034

Schneider, D., Mertes, C., & Wascher, E. (2016). The time course of visuo-spatial working memory updating revealed by a retro-cuing paradigm. Sci Rep, 6, 21442. doi:10.1038/srep21442

Sligte, I. G., Scholte, H. S., & Lamme, V. A. (2008). Are there multiple visual short-term memory stores? PLoS One, 3(2), e1699. doi:10.1371/journal.pone.0001699

Snyder, A. C., & Foxe, J. J. (2010). Anticipatory attentional suppression of visual features indexed by oscillatory alpha-band power increases: a high-density electrical mapping study. J Neurosci, 30(11), 4024–4032. doi:10.1523/JNEUROSCI.5684-09.2010

Stokes, M. G. (2015). ’Activity-silent’ working memory in prefrontal cortex: a dynamic coding framework. Trends Cogn Sci, 19(7), 394–405. doi:10.1016/j.tics.2015.05.004

Vandenbroucke, A. R., Sligte, I. G., & Lamme, V. A. (2011). Manipulations of attention dissociate fragile visual short-term memory from visual working memory. Neuropsychologia, 49(6), 1559–1568. doi:10.1016/j.neuropsychologia.2010.12.044

Wang, B., & Theeuwes, J. (2018). How to inhibit a distractor location? Statistical learning versus active, top-down suppression. Atten Percept Psychophys, 80(4), 860–870. doi:10.3758/s13414-018-1493-z

Whitney, P., Arnett, P. A., Driver, A., & Budd, D. (2001). Measuring central executive functioning: what’s in a reading span? Brain Cogn, 45(1), 1–14. doi:10.1006/brcg.2000.1243

Williams, M., Hong, S. W., Kang, M. S., Carlisle, N. B., & Woodman, G. F. (2013). The benefit of forgetting. Psychon Bull Rev, 20(2), 348–355. doi:10.3758/s13423-012-0354-3

Wolff, M. J., Jochim, J., Akyurek, E. G., & Stokes, M. G. (2017). Dynamic hidden states underlying working-memory-guided behavior. Nat Neurosci, 20(6), 864–871. doi:10.1038/nn.4546

Woodman, G. F., & Luck, S. J. (2003). Serial deployment of attention during visual search. J Exp Psychol Hum Percept Perform, 29(1), 121–138.

