## supplementary figure S1 for "Hemispheric asymmetries in posterior alpha power reflect the selection and inhibition of spatial context information in working memory" (from another version of this preprint: version 2, 2019-07-09)

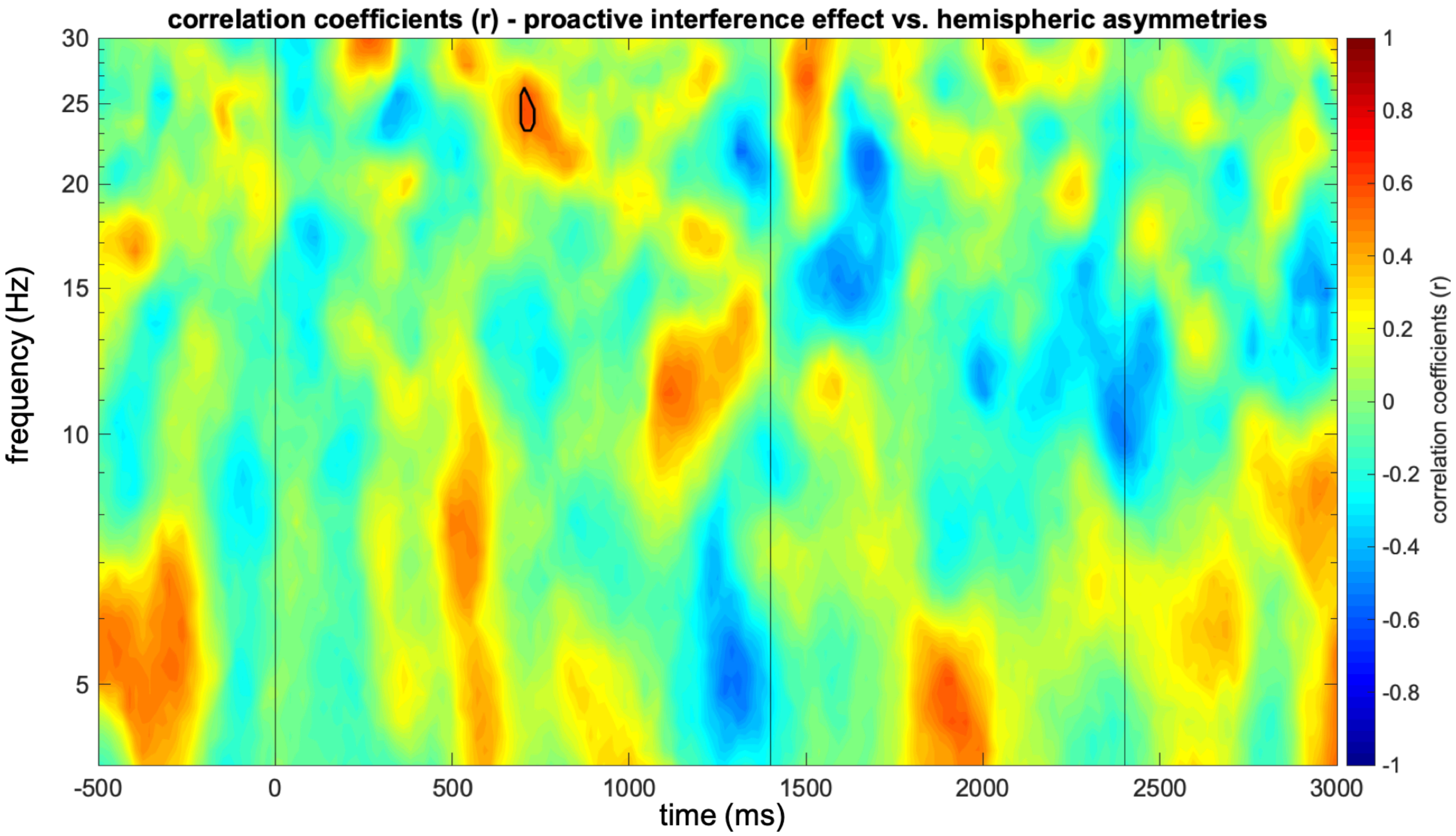


**Supplementary figure S1.** *Between-subject correlations for posterior hemispheric asymmetries. The plots show the correlations coefficients for each time-frequency point based on the difference of hemispheric asymmetries (PO7/8, PO3/4, P7/8, P5/6) between the distractor and target lateral conditions and the proactive interference effect on RT level. Vertical lines indicate the onsets of the memory array, retro-cue and memory probe. Results indicate that there was no consistent relation between oscillatory correlates of spatial retroactive attentional orienting and the ability to control for behavioral consequences of proactive interference in the current dataset.*
